# Large-scale pan-cancer analysis reveals broad prognostic association between TGF-β ligands, not Hedgehog, and GLI1/2 expression in tumors

**DOI:** 10.1101/728949

**Authors:** Aurélien de Reyniès, Delphine Javelaud, Nabila Elarouci, Véronique Marsaud, Cristèle Gilbert, Alain Mauviel

## Abstract

*GLI1* expression is broadly accepted as a marker of Hedgehog pathway activation in tumors. Efficacy of Hedgehog inhibitors is essentially limited to tumors bearing activating mutations of the pathway. GLI2, a critical Hedgehog effector, is necessary for *GLI1* expression and is a direct transcriptional target of TGF-β/SMAD signaling. We examined the expression correlations of GLI1/2 with *TGFB* and *HH* genes in 152 distinct transcriptome datasets totaling over 23,500 patients and representing 37 types of neoplasms. Their prognostic value was measured in over 15,000 clinically annotated tumor samples from 26 tumor types. In most tumor types, *GLI1* and *GLI2* follow a similar pattern of expression and are equally correlated with *HH* and *TGFB* genes. However, *GLI1*/*2* broadly share prognostic value with *TGFB* genes and a mesenchymal/EMT signature, not with *HH* genes. Our results provide a likely explanation for the frequent failure of anti-Hedgehog therapies in tumors, as they suggest a key role for TGF-β, not Hedgehog, ligands, in tumors with elevated *GLI1/2*-expression.

## Introduction

Elevated expression and nuclear localization of GLI1, reminiscent of Hedgehog (HH) pathway activation, has been reported in a wide variety of tumor types [1]. Although extensive experimental evidence exists for a pro-tumorigenic and pro-metastatic role of GLI1, efficacy of HH inhibitors is restricted to a handful of cancers with genetic activation of upstream components of the pathway [2, 3]. The sole FDA-approved indication for HH inhibitors is advanced cutaneous basal cell carcinoma [4]. Understanding their lack of efficacy in other tumors despite high GLI1 expression remains a challenge [5, 6].

Members of the HH family of growth factors, Sonic (SHH), Indian (IHH) and Desert (DHH) control tissue patterning, limb and skeletal polarity during embryonic life, and broadly contribute to tissue homeostasis and repair processes during adulthood, by controlling cell proliferation, migration, as well as stem cell maintenance and self-renewal [1, 7]. Ligand binding to the 12-transmembrane receptor PATCHED-1 (PTCH1) allows activation of the 7-transmembrane G-coupled receptor Smoothened (SMO), and HH signal transduction proceeds towards activation and nuclear accumulation of GLI transcription factors with activator or repressor functions dependent upon proteolytic cleavage [8]. *GLI1* is a prototypic HH target gene and its expression is widely considered a read-out of HH pathway activation. GLI2 is the primary substrate and effector of the pathway that largely contributes to *GLI1* induction by HH, as well as that of other HH target genes in cooperation with GLI1 [9, 10]. GLI3 has weak transcriptional activity and is considered an inhibitor of HH activity [8].

Ligand-independent HH pathway activation as a result of mutations in genes encoding upstream pathway components, such as loss-of-function mutations in *PTCH1* or *SUFU* (suppressor of fused) and activating mutations of *SM0* are rare in cancers and only found in cutaneous basal cell carcinoma, rhabdomyosarcoma, and medulloblastoma, cancers which exhibit notable therapeutic response to HH inhibitors [11].

Clinical studies on HH inhibitors have often relied on increased expression or nuclear localization of GLI1 as a marker of HH pathway activation. It may be argued that in a number of cases, HH activation was stated empirically, with no direct evidence of active upstream HH signaling. Similarly, studies targeting GLI1/2 expression or function in tumor cells *in vitro* or in mouse models of cancer have shown remarkable anti-tumor efficacy and concluded to a pathogenic role for the HH pathway, although most studies targeted its main downstream effectors, not the pathway itself. Thus, while there is little doubt that GLI transcription factors contribute substantially to cancer progression, direct evidence that would link GLI activity in a given tumor setting to HH ligands activating their receptors is often missing. This may explain the overall lack of anti-tumor therapeutic efficacy of HH inhibitors, most of which target SMO activity [1].

We identified TGF-β as a powerful inducer of GLI2 and GLI1 expression as well as GLI-dependent transcription, independent from SMO activity [12, 13]. We established a role for both TGF-β signaling and GLI2 in driving melanoma invasion and metastasis, that could be targeted with TGF-β receptor inhibitors or by knocking down GLI2 expression [14–16]. Similar observations have since been reported for other tumor types, including ovarian and oral squamous cell carcinomas, that link GLI2 and TGF-β expression to tumor aggressiveness [17–19].

Herein, we hypothesized that the lack of efficacy of SMO antagonists in numerous tumors occurs because high *GLI1* expression and activity may not be linked to HH, but rather to TGF-β, ligand expression, taken as surrogates for HH and TGF-β signaling in tumors, irrespective of the cellular compartment. We compiled data from publicly available gene expression datasets from over 23,500 cancer patients, of which over 15,000 with survival annotations, well above those available from The Cancer Genome Atlas (TCGA). While *GLI1*/*2* expression is correlated with both *HH* and *TGFB* expression, their prognostic value is tightly correlated with that of *TGFB*, not *HH*. High *GLI1*/*2* and *TGFB* expression, associated with a mesenchymal/EMT signature, often represent parallel markers of poor clinical outcome. Inversely, high expression of *HH* is mostly associated with increased survival.

## Results

### Pan cancer correlation between GLI1, GLI2, TGFB, and HH genes

We hypothesized that the correlation between *GLI1* and *GLI2* expression with HH (*SHH*, *IHH*, *DHH*) and TGF-ß ligands (*TGFB1*, *TGFB2*, *TGFB3*) transcript levels represent adequate surrogates for the respective pathogenic implication of HH and TGF-ß ligands in GLI1/2 expression and activity in tumors. Pan-cancer analysis of the correlation of *GLI1* and *GLI2* with 19540 genes expressed in 30 tumor types revealed that *GLI1* and *GLI2* are mutually the most correlated genes (Figure 1A), and that all *HH* and *TGFB* genes are in the top 20 percentiles of most correlated genes with *GLI1* and *GLI2*, with one exception. Expression of at least one of the *TGFB* genes was more closely related than that of any *HH* genes to both *GLI1* and *GLI2* expression.

**Figure 1.**
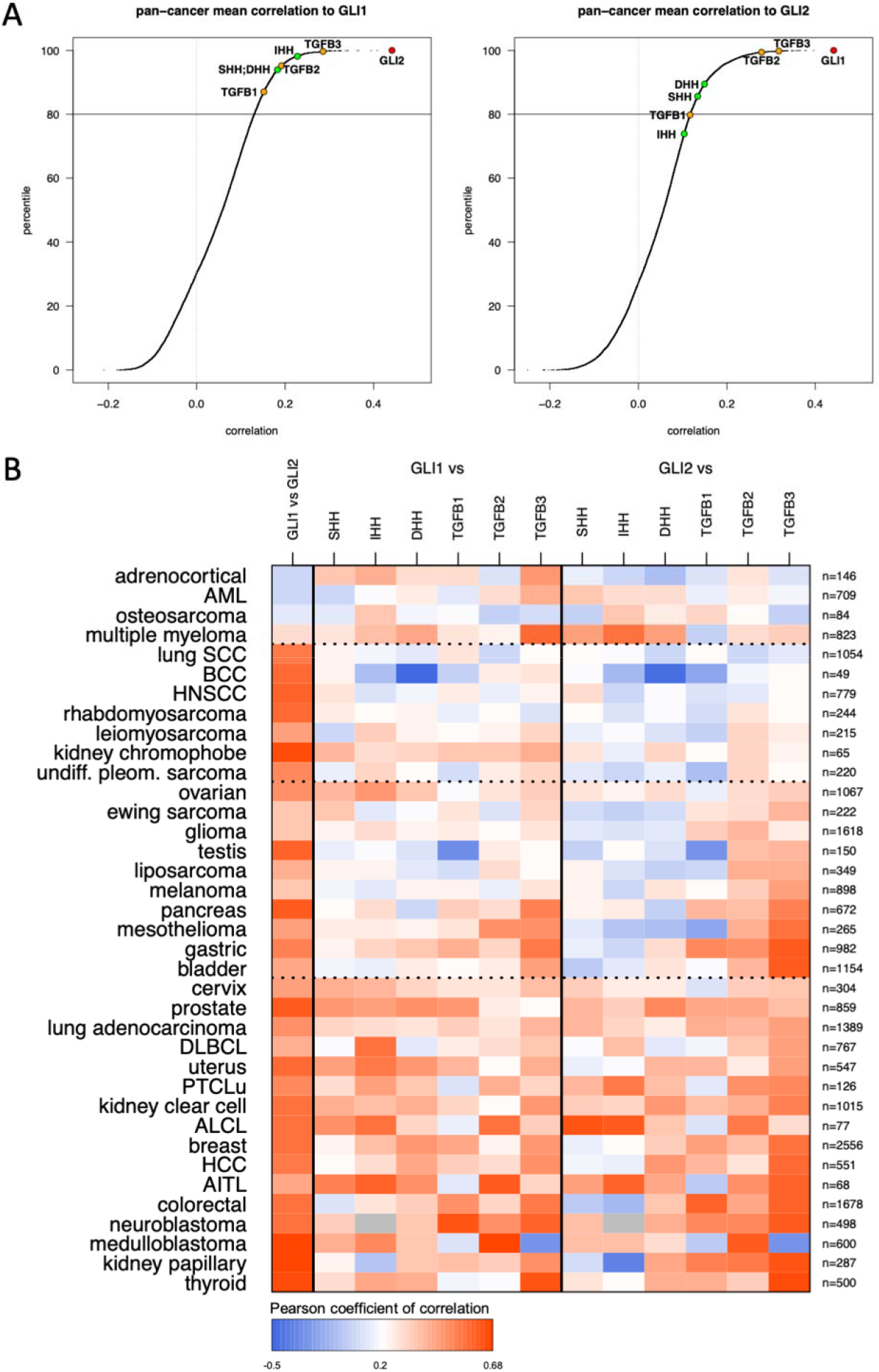
Pan-cancer expression correlations with *GLI1* and *GLI2*. **A**. Data from 152 expression public datasets from 37 cancer types, spanning over 23.500 patients were sorted and the correlation of 19540 genes (expressed in at least 30 tumor types) with that of *GLI1* (left panel) and *GLI2* (right panel) was calculated. The respective position of *HH* and *TGFB* genes is indicated. **B**. Heatmap representation of the expression correlation between *GLI1* and *GLI2* with each other and between *HH* and *TGFB* genes in 37 cancer types. Corresponding numerical values are provided in Supplementary Table 2. The number of patients for each neoplasm is indicated.

Positive correlation (arbitrary threshold: r>0.25) between the *GLI1* and *GLI2* genes was observed in 33/37 tumor types (Figure 1B), representing 92.5% (21825/23587) of patients. Most tumor types exhibited similar correlation values between *GLI1* expression and that of at least one of either *HH* or *TGFB* genes (Figure 1B). The correlation pattern between *HH* and *TGFB* genes with *GLI2* (Figure 1B) was similar to that with *GLI1*, yet tumors with high *GLI2*/*TGFB* correlation could be discriminated into two subgroups: one exhibiting low *GLI2*/*HH* correlation (tumor types from ovarian down to bladder), the other exhibiting high *GLI2*/*HH* correlation (tumor types from cervix down to thyroid). We did not identify a single neoplasm for which *GLI1*/*2* expression was correlated with that of *HH* genes without a simultaneous correlation with that of at least one of the *TGFB* genes.

### Expression of GLI1, GLI2, HH and TGFB genes differentially associates with key oncogenic signatures

Cell cycle progression, acquisition of a mesenchymal phenotype through epithelial-to-mesenchymal transition (EMT), and cell stemness are cellular traits considered hallmarks of cancer progression [20], to which both the HH and TGF-ß pathways are linked [2, 3]. For each of the 37 tumor types, we generated a multivariate linear model based on the expression of the eight genes of interest taken together (three *HH* and three *TGFB* genes, *GLI1* and *GLI2*) to determine whether it may be predictive of these metagenes. To assess the goodness-of-fit of these models, correlations between the predicted values and observed values for each metagene were calculated in each tumor type. As shown in Figure 2A, strong correlations were observed in most tumor types for the mesenchymal/EMT and cancer cell stemness metagenes albeit to a lesser extent, while it was seldom observed with the cell cycle metagene.

**Figure 2.**
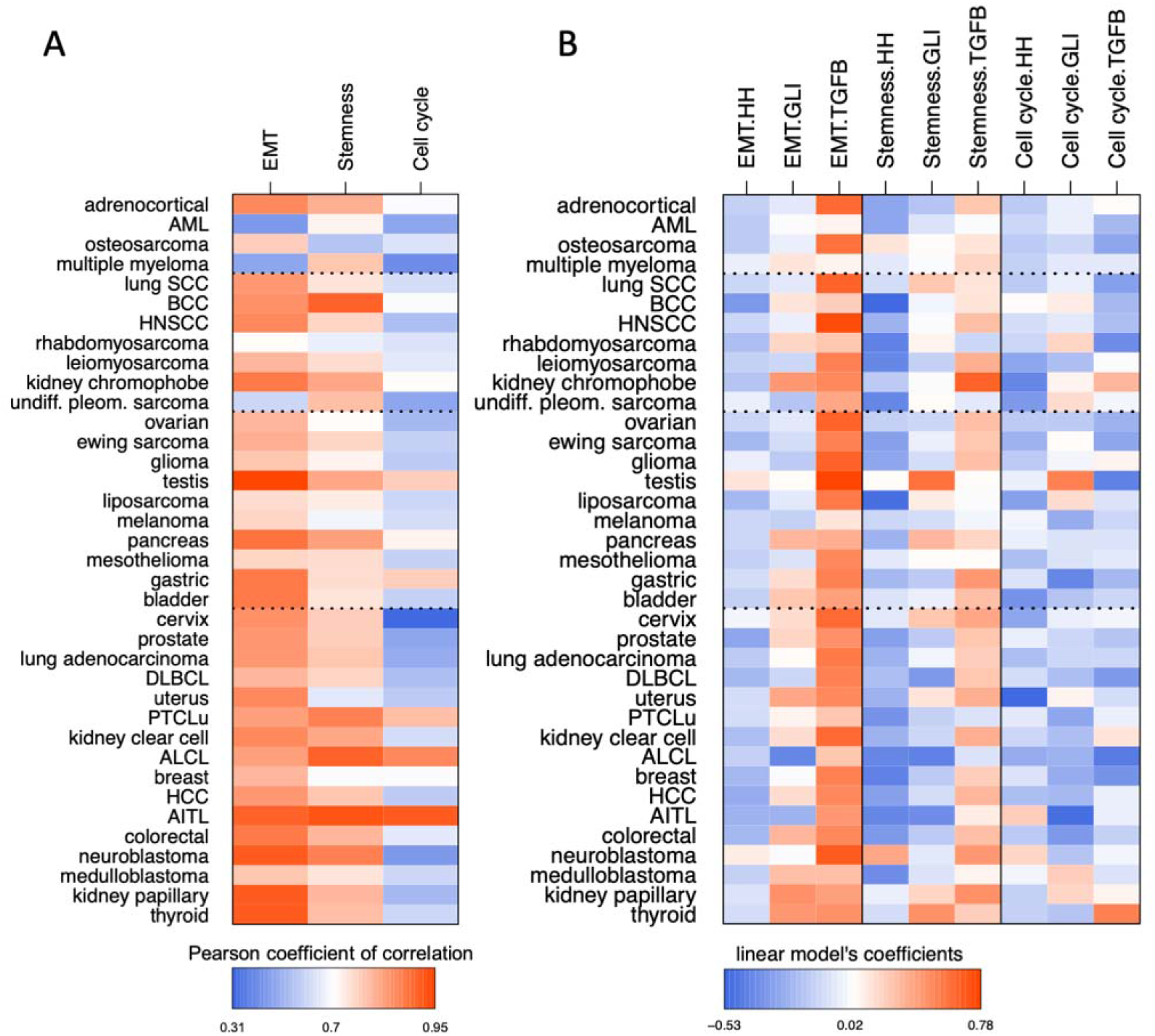
Multivariate linear prediction models of metagene signatures for select major oncogenic traits (mesenchymal/EMT, stemness, cell cycle). **A**. Heatmap representation of the correlations between predicted and observed values of metagenes in 37 tumor types, taking *GLI1*/*GLI2*, the three *HH* and three *TGFB* genes together. Corresponding numerical values are provided in Supplementary Table S4B. **B**. Heatmap representation of the coefficients from the linear models for each of the 3 metagenes in each tumor type, based on the combined expression of either *GLI1* and *GLI2,* or the three *HH* or *TGFB* genes. Corresponding numerical values are provided in Supplementary Table S4B.

A simplified multivariate linear model using compounded expression of either *GLI1*/*2*, the *HH* or *TGFB* genes was next calculated for each metagene. Coefficients for these three predictive variables within each model, presented in Figure 2B, demonstrate the dominant role of *TGFB* gene expression, followed by that of *GLI1/2*, not *HH*, in predicting mesenchymal and cancer cell stemness metagene expression in a broad array of tumor types. None of them was associated with the cell cycle metagene. Data for each *GLI*, *HH* and *TGFB* gene taken individually in each model are provided in Supplementary Figure S1.

### Pan-cancer prognostic values associated with GLI1/2, HH and TGFB transcript levels and select oncogenic signatures

Univariate Cox survival analysis from over 15,000 clinically annotated tumor samples brought critical information (Figure 3A). At odds with a generalized assumption in the clinical setting that HH signaling is deleterious, expression of all *HH* genes in tumors was associated with good prognosis (H.R.<1, green color). Not a single tumor type was found for which expression of any *HH* gene was associated with morbidity (H.R.>1, red color) without a parallel pejorative prognostic value for at least one *TGFB* gene and either *GLI1*, or *GLI2*. Also, for each occurrence when high *GLI1* or *GLI2* expression was of bad prognosis, the same held true for at least one of the *TGFB* genes. On the other hand, when either *GLI1* and *GLI2* expression were of good prognosis, the same applied for at least one of the *TGFB* genes. Noticeably, in bladder, colorectal and kidney papillary carcinoma, *GLI2* (and *GLI1*) expression shared bad prognostic value with that of *TGFB genes*, while high *HH* expression was associated with a positive outcome. As expected, the three metagenes were largely of bad prognosis, with mesenchymal/EMT and cancer cell stemness metagenes exhibiting a largely overlapping, yet cancer type-specific pattern of prognostic significance, while the cell cycle metagene was almost universally associated with poor outcome.

**Figure 3.**
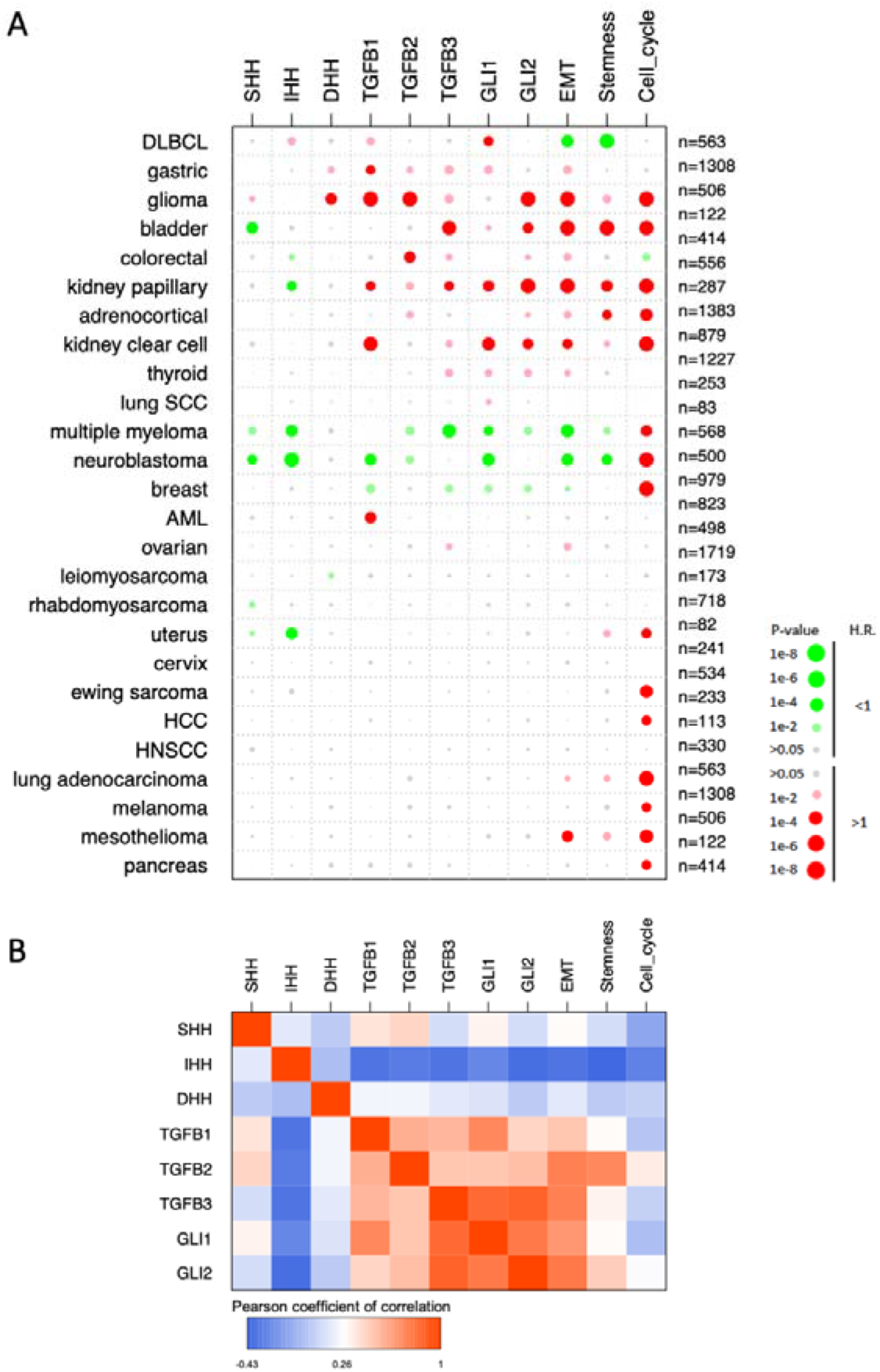
Prognostic value associated with *GLI(1/2)*, *HH(S/I/D)* and *TGFB(1/2/3)* genes and the mesenchymal/EMT, cell stemness and cell cycle metagenes in human cancers. **A**. Data were derived from a meta-analysis for the univariate prognostic value for overall survival in 26 types of human cancers for which sufficient events were available. Bigger circles represent lower *p* values. Marked colors represent *p* value below 0.001, dull colors represent *p* values below 0.05. Green and red colors represent H.R. below and above 1, respectively (see escale bar)**. B**. Correlations between pan-cancer profiles of prognostic scores of *GLI(1/2)*, *HH(S/I/D)* and *TGFB(1/2/3)* genes and the mesenchymal/EMT, cell stemness and cell cycle metagenes. Prognostic scores are defined as the log2(H.R.) if the related *p*-value is below 0.05, or 0 otherwise.

In breast cancer, *HH* expression had no prognostic value while *GLI1*/*2* and *TGFB* expression were associated with better survival, together with the mesenchymal/EMT and cell stemness metagenes. These results are at odds with most neoplasms where *GLI1*/*2* and *TGFB* genes share pejorative prognostic value. To understand this discrepancy, a large cohort of breast cancer patients’ data [21] was further analyzed. Intra-dataset z-score *GLI*(1/2), *TGFB*(1/2/3) and *HH*(S/I/D) and metagene expression values were sorted according to increasing *GLI2* expression, then aligned to the molecular subtypes. High *GLI1/2/TGFB* and mesenchymal/EMT metagene expression was associated with the normal-like subgroup of tumors with better prognosis, and inversely correlated with luminal-type tumors of poor prognosis (Supplementary Figure S2). A hypothesis may be that the mesenchymal/EMT metagene, which follows *GLI2* and *TGFB* expression pattern, may not be representative of actual EMT in normal-like breast tumors, but rather represents that of fibroblastic heterogeneity found in breast tumors [22].

For each cancer type, we next defined a prognostic score as the log2(H.R.) if the related p-value was below 0.05, or 0 otherwise. Correlations between the prognostic scores for *GLI(1/2), TGFB(1/2/3), (S/I/D)HH* and oncogenic function metagenes were calculated. The results, presented in Figure 3B, overwhelmingly demonstrate that *GLI*(1/2), *TGFB*(1/2/3), Mesenchymal/EMT and stemness metagenes have highly correlated pan-cancer prognostic profiles, with no or modest correlation to that of *HH*(S/I/D). Little or no correlation between *GLI(1/2), TGFB(1/2/3), (S/I/D)HH* was found with the cell cycle metagene H.R, consistent with the expression correlation data from Figure 2.

## Discussion

HH inhibitors have failed to fulfill expectations placed on them as powerful anti-cancer drugs. Despite widespread expression of *GLI1* in tumors, considered to be a marker of HH activation, HH inhibitors have solely been granted FDA approval for the treatment of advanced basal cell carcinoma of the skin, a type of tumor that bears activating mutations of the HH pathway. Clinical trials on other solid (or non-solid) tumors have overall failed. Based on our earlier work that identified TGF-ß as a potent transcriptional inducer of GLI2, consequently leading to SMO-independent induction of GLI1 [12, 13], we hypothesized that *GLI1* expression in tumors may be driven by TGF-ß, not HH, which would explain the lack of efficacy of anti-HH approaches. Large-scale datamining from gene expression datasets representing 23,500 patients and 37 types of neoplasms addressed the issue of whether *GLI1*/*2* expression in tumors warrants therapeutic approaches targeting upstream HH signaling. While significant correlation was found between the expression of *GLI1*/*2* and *HH* genes in tumors, correlation with that of *TGFB* genes was as strong. Thus, not only *GLI1* is not a relevant marker to predict HH pathway activity, but neither *GLI1* nor *GLI2* expression in tumors discriminate between HH and TGF-ß ligands as potential upstream inducers of their expression.

Strikingly, the prognostic value of *GLI1*/*2* expression in tumors was mostly at odds with that of *HH* genes, while paralleling that of one or more *TGFB* genes, as well as that of mesenchymal/EMT and cell stemness metagenes. Contrary to the pejorative prognosis associated with high *GLI1*/*2*/*TGFB* gene expression, high *HH* expression was mostly associated with good prognosis. In the rare occurrences when *HH* expression was associated with poor outcome, at least one *TGFB* gene shared the prognosis. This is schematized in Figure 4.

**Figure 4.**
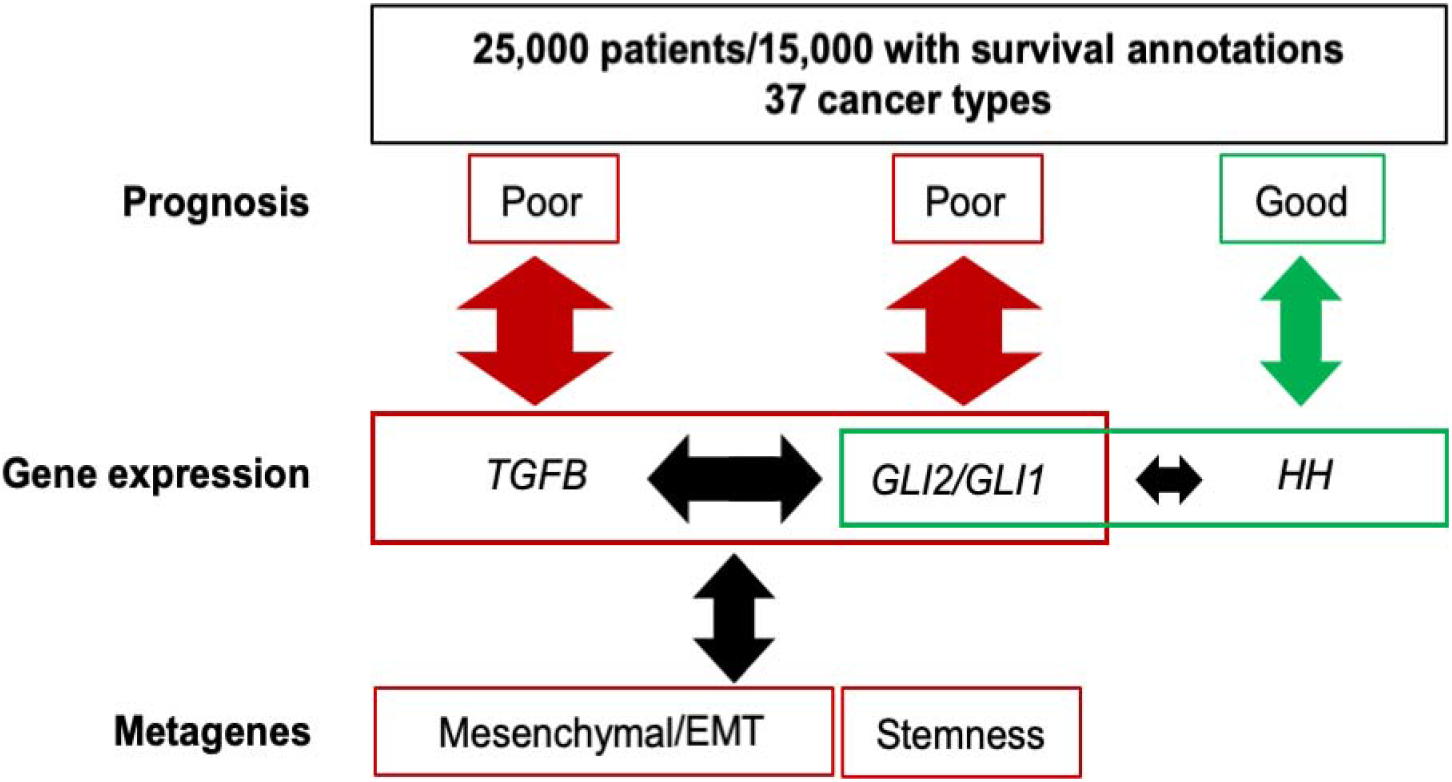
Schematic summary. Pan cancer analysis of gene expression and survival data demonstrates shared poor prognostic value for *TGFB*, *GLI1/2*, together with mesenchymal/EMT and Stemness signatures, not *HH*. Arrow size illustrates the relative correlations between genes (TGFB, GLI2/GLI1 or HH), prognosis and metagene signatures.

Our broad pan-cancer analysis herein indicates that GLI1/2 functions parallel or match those of TGF-ß ligands, not HH, as identified both in linear models of oncogenic functions and in prognostic analyses of tumors. Multivariate analysis demonstrated the dominant role of *TGFB*, followed by that of *GLI1/2*, not *HH*, in predicting mesenchymal/EMT and cell stemness metagene expression in a broad array of tumor types. These data fit a mechanistic model whereby TGF-ß controls *GLI2* expression which, in turn and depending upon context, allows for *GLI1* expression in either a HH-dependent or -independent fashion, as proposed previously [23].

Our data hint that HH ligand-driven signaling in tumors leading to *GLI1* expression without an overlap and contribution of the TGF-ß pathway is not only a rare event but is also unlikely to be pathogenic. The divergence between the prognostic value of *GLI1*/*2* expression and that of *HH* ligands indicates that there is no sensible justification for targeting HH for cancer treatment if *GLI1* expression is used as a prognostic variable. Patients’ selection based upon an inadequate marker of HH pathway activation may therefore contribute to the lack of clinical efficacy of SMO antagonists in various neoplasms. The uniqueness of the cell cycle metagene prognostic value is independent from *GLI1*/*2*, *TGFB* and *HH* gene signatures. Together with its broad pan-cancer pejorative prognostic value distinct from that of *TGFB*/*GLI1*/*2*/EMT/Stemness, suggests a potential therapeutic benefit for the combination of cytostatic drugs together with anti TGF-ß/GLI inhibitors.

## Material and Methods

### Transcriptome series

A set of 135 transcriptome series related to 37 cancer types was collected from public repositories (ArrayExpress, GEO, TCGA). Series that included multiple cancer types were split accordingly, yielding to a total of 152 distinct transcriptome datasets representing over 23,500 patients. Details and accession numbers are provided in Supplementary Table S1.

### Pre-treatment, normalization

Datasets based on Affymetrix microrrays were normalized independently using the justRMA function from the Affy R package (with default parameter). For TCGA RNA-Seq datasets, raw counts were normalized using the upper quartile method [24]. Datasets from other sources were used as furnished, after log2 transformation of expression values. All probe sets data were aggregated by HUGO Gene Symbol.

### Correlation analyses

Transcripts with available measures in at least 30 cancer types were selected for further analyses (n=19,540). The correlation between their expression and that of *GLI1* and *GLI2* was calculated independently in each of the 152 datasets, yielding 152 correlation matrices (dimension: 19,540×2). Correlations were then averaged across all 152 dataset matrices, yielding a unique (19,540×2) matrix used to plot Fig. 1A. The 152 matrices were also averaged per cancer type, yielding 37 sub-matrices that were reduced to the 8 genes of interest (*GLI1*, *GLI2*, *TGFB*(1/2/3) and *HH*(S/I/D), Supplementary Table S2) and used to plot Fig. 1B.

### Metagene calculation

Three published gene signatures corresponding to critically important oncogenic activities were selected: (i) mesenchymal-EMT [25], (ii) Stemness [26]; (iii): Cell Cycle: https://www.genome.jp/dbget-bin/www_bget?pathway+hsa04110. Gene content for each signature is listed in Supplementary Table S3. For each of these signatures and each dataset, the average zero-centered expression of the genes, both measured and included in the signature, was calculated for each sample.

### Linear models

Within each dataset, based upon the metagene values for each of the three oncogenic signatures, the lm function from the stats R package was used to perform a linear regression of the metagene variable, using *GLI1*, *GLI2*, *SHH*, *IHH*, *DHH*, *TGFB1*, *TGFB2* and *TGFB3* expression as predictive variables. To allow for inter-datasets and inter-variables comparisons, all variables were z-scored within each dataset before linear modeling (common unit=standard deviation). For each model in each dataset, correlations between predicted values and observed metagene values (z-scores) was recorded and averaged across datasets by cancer type, then represented as heatmaps. For each linear model in each dataset, coefficients of the predictive variables were recorded and averaged across datasets by cancer type and metagene. Numerical values are provided per series and per cancer type (Supplementary Tables S4A and S4B, respectively).

### Survival analyses

Univariate Cox models of overall survival in 26 tumor types with cohorts of more than 50 patients and at least 10 death events were calculated using the survival R package. Aggregation of Hazard Ratios (H.R.) and related confidence intervals across datasets of a given cancer type for a given variable were calculated using the meta R package. All genes and metagenes were z-scored intra-dataset prior to modeling, to allow for inter-datasets comparisons (common unit=standard deviation). Numerical values are provided per series and per cancer type (Supplementary Tables S5A and S5B, respectively).

## Acknowledgments

This work was supported by the French national program Cartes d**’**Identité des Tumeurs® (CIT) funded and developed by the Ligue Nationale Contre le Cancer, grants from Ligue Nationale Contre le Cancer (Equipe Labellisée LIGUE-2016), Institut National du Cancer (INCa MELA13-002) and Donation Henriette et Emile Goutière (to A.M), Fondation ARC (to D.J.), and institutional funding from Institut National de la Santé et de la Recherche Médicale, Centre National de la Recherche Scientifique, Institut Curie, and Université Paris-Sud.

## Disclosure statement

The authors have no conflict of interest to declare.

## Supporting information

**Figure S1**: Heatmap representation of the coefficients from the linear models generated for each of the 3 metagenes in each tumor type, based on the expression of *GLI1*, *GLI2*, *TGFB1*, *TGFB2*, *TGFB3*, *SHH*, *IDH*, *DHH*.

**Figure S2**: Alignment of *GLI1*, *GLI2*, *SHH*, *DHH*, *IHH*, *TGFB1*, *TGFB2* and *TGFB3* as well as mesenchymal/EMT, Stemness and Cell cycle metagene expression profiles with molecular subtypes of breast carcinomas

**Table S1**: Public datasets

**Table S2**: Correlations (pearson coef.) per cancer type

**Table S3**: Genesets used for metagene calculation

**Table S4**: Multivariate linear models, A: intra series; B: aggregation per cancer type

**Table S5**: Univariate Cox models, A: intra series; B: aggregation per cancer type

