## Supplementary Figs 1 and 2 for "Large-scale pan-cancer analysis reveals broad prognostic association between TGF-β ligands, not Hedgehog, and GLI1/2 expression in tumors"

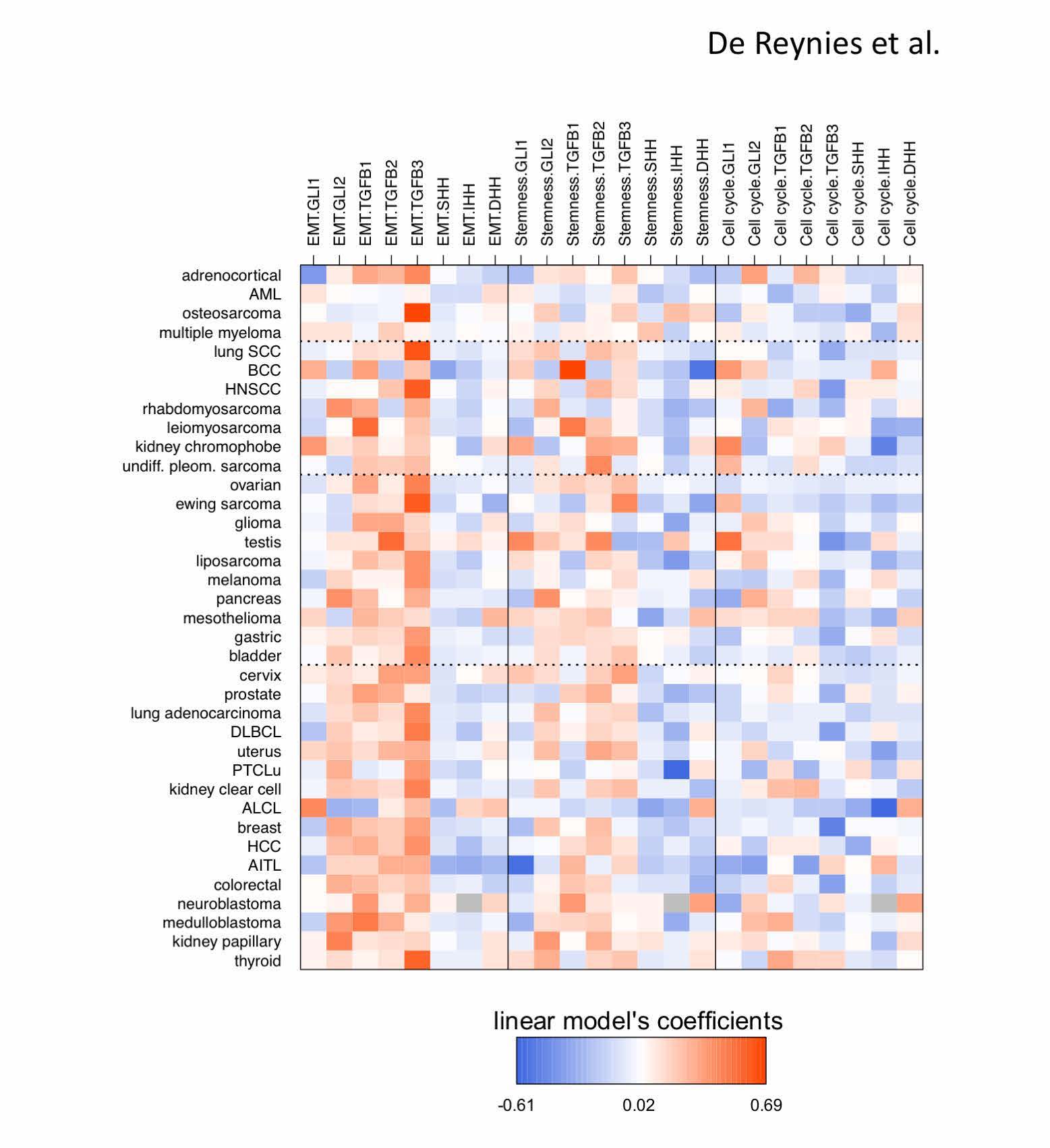

**Supplementary Figure S1.** Heatmap representation of the coefficients from the linear models generated for each of the 3 metagenes in each tumor type, based on the expression of *GLI1*, *GLI2*, *TGFB1*, *TGFB2*, *TGFB3*, *SHH*, *IDH*, *DHH*. The corresponding numerical values are provided in Supplementary Table S4B.

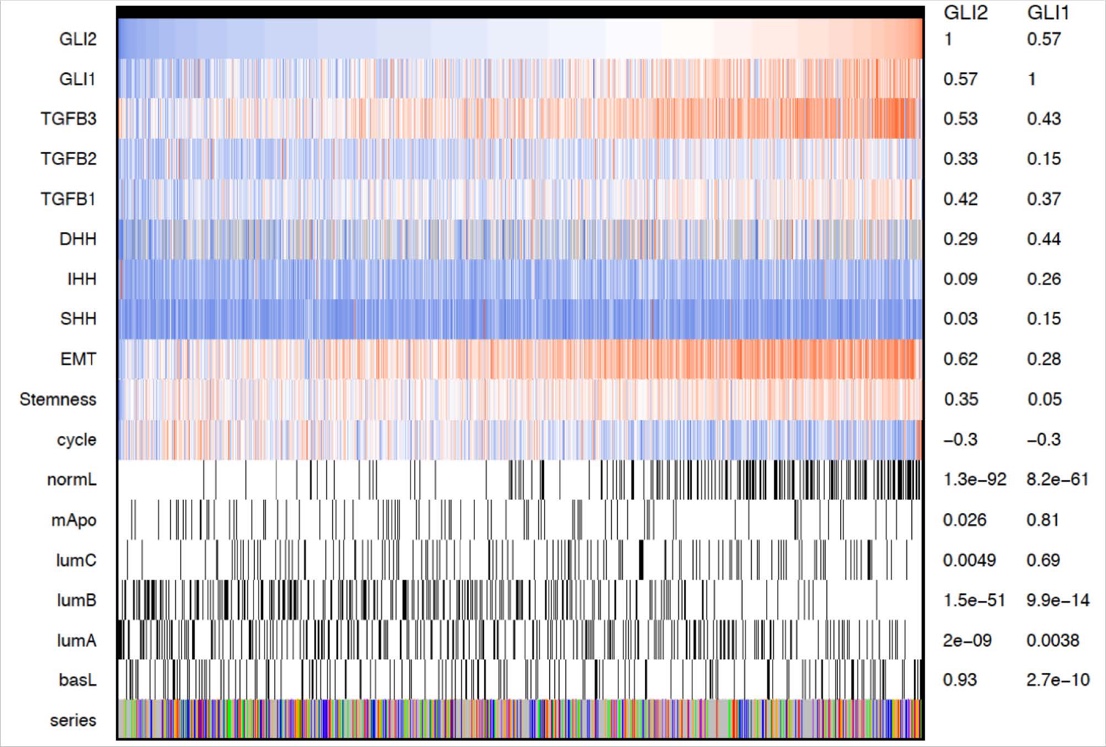

**Supplementary Figure S2.** Alignment of *GLI1*, *GLI2*, *SHH*, *DHH*, *IHH*, *TGFB1*, *TGFB2* and *TGFB3* as well as mesenchymal/EMT, Stemness and Cell cycle metagene expression profiles with molecular subtypes of breast carcinomas (6 series, 2556 patients).
